# Detection of Frog virus 3 via the system integrating RPA-CRISPR/Cas12a-SPM with deep learning

**DOI:** 10.1101/2022.08.22.504785

**Authors:** Zhengyang Lei, Lijin Lian, Likun Zhang, Jiazhang Wei, Changyue Liu, Hong Liu, Ying Liu, Zhicheng Du, Xi Yuan, Xiaoyun Zhong, Ijaz Gul, Haihui Zhang, Chuhui Wang, Shiyao Zhai, Vijay Pandey, Canyang Zhang, Xinhui Xing, Lin Deng, Dongmei Yu, Qian He, Peiwu Qin

## Abstract

Frog virus 3 (FV3, genera *Ranavirus*, family *Iridoviridae*), a double-stranded DNA virus, results in irreparable damage to biodiversity and significant economic losses to aquaculture. Although the existing FV3 detection methods are of high sensitivity and specificity, the complex procedure and requirement of expensive instruments limit their practical implantation. Herein, we develop a fast, easy-to-implement, highly sensitive, and point-of-care (POC) detection system for FV3. Combining recombinase polymerase amplification (RPA) and CRISPR/Cas12a, we achieve a limit of detection (LoD) of 100 aM (60.2 copies/μL) by optimizing RPA primers and CRISPR RNAs (crRNAs). For POC detection, we build a smartphone microscopy (SPM) and achieve an LoD of 10 aM within 40 minutes. Four positive animal-derived samples with a quantitation cycle (Cq) value of quantitative PCR (qPCR) in the range of 13 to 32 are detectable by the proposed system. In addition, we deploy deep learning models for binary classification (positive or negative samples) and multiclass classification (different concentrations of FV3 and negative samples), achieving 100% and 98.75% accuracy, respectively. Without temperature regulation and expensive equipment, RPA-CRISPR/Cas12a combined with a smartphone readout and artificial intelligence (AI) assisted classification shows great potential for FV3 detection. This integrated system holds great promise for POC detection of aquatic DNA pathogens.

**Highlights:**

1. An integrated DNA detection system is developed by combining RPA, CRISPR/Cas12a, smartphone microscopy, and deep learning.
2. The LoD of frog virus 3 is 10 aM within 40 min.
3. The detection system shows good performance on animal-derived samples.

## 1. Introduction

Ranaviruses, including the type species *frog virus 3* (FV3), are double-stranded DNA (dsDNA) viruses within the family *Iridoviridae* (Jancovich et al. 2010; Robert and Gregory 2012). Diseases caused by ranaviruses are a major concern for biodiversity and aquaculture (Chen and Robert 2011; Price et al. 2014; Whittington et al. 2010; Youker-Smith et al. 2018; Yu et al. 2021). FV3 infections induce hematopoietic necrosis in the bone marrow, lesions in the skin and hematopoietic tissue, and necrosis of lymphoid tissue and mucosal epithelium, which results in the death of the host within a few days to several weeks (Forzan et al. 2017a; Morrison et al. 2014). FV3 causes 90 - 100% mortality in a wide range of amphibian species, which significantly contributes to global amphibian decline (Hossainey et al. 2021; Hoverman et al. 2011; Miller et al. 2007). However, there are no effective preventative vaccines on the market (Chinchar et al. 2017). Therefore, a rapid and accurate detection system is urgently required to prevent the spread of FV3.

There is no single Gold Standard test for FV3 detection (Miller et al.). The traditional detection methods include histopathology examination, virus isolation, enzyme-linked immunosorbent assay (ELISA), polymerase chain reaction (PCR), and quantitative real-time PCR (qPCR) (Fong and Lipp 2005). The major capsid (MCP) gene is highly conserved and usually selected as the target for ranaviruses detection by PCR. The limit of detection (LoD) by PCR and qPCR for FV3 has not been reported. For our reference, targeting the MCP gene, the LoD for *largemouth bass virus* (LMBV) by PCR and qPCR is 1000 copies/μL and 5.8 copies/μL in more than 60 min (Guo et al. 2022). However, these common methods are complicated, costly, and time-consuming for point-of-care (POC) detection.

Recently, clustered regularly interspaced short palindromic repeats (CRISPR) and CRISPR-associated (Cas) protein have enticed substantial attention for nucleic acid detection and have shown promising results (Bao et al. 2021; Broughton et al. 2020; Chen et al. 2018a; Gootenberg et al. 2017; Kellner et al. 2019; Mukama et al. 2020; Schwank et al. 2013; Yin et al. 2021). Cas12a protein guided by CRISPR RNA (crRNA) binds and cleaves target DNA, unleashing nonspecific single-stranded DNA (ssDNA) trans-cleavage activity (Liang et al. 2019; Schunder et al. 2013; Zetsche et al. 2015), which makes it suitable for nucleic acid detection (Chen et al. 2018b; He et al. 2020a; Li et al. 2018; Qin et al. 2019). We previously developed a CRISPR/Cas12a-based automated, integrated, and inexpensive detection system for *African swine fever virus* (ASFV). A detection limit of 1 pM is achieved in 2 hours without amplification (He et al. 2020a). To improve the sensitivity for trace viral DNA detection, the CRISPR/Cas12a system can be combined with recombinase polymerase amplification (RPA) or loop-mediated isothermal amplification (LAMP), which has been used for detecting multiple kinds of pathogens, such as *Plasmodium falciparum, Plasmodium vivax, scale drop disease virus*, ASFV, and SARS-CoV-2 (Bai et al. 2019; Lee et al. 2020; Mayuramart et al. 2021; Sukonta et al. 2022). However, LAMP is limited in use because the design of the primers is complicated (Mh and Nck 2021). Therefore, we combine RPA and CRISPR/Cas12a for FV3 detection that remains unexplored to the best of our knowledge.

For POC detection, the handheld fluorimeter, lateral flow strips, or smartphone microscopy (SPM) is developed for the results readout (Fozouni et al. 2021; Kumar et al. 2021; Lee et al. 2020). SPM uses a camera to capture images, which can be uploaded to mobile applications for fast data analysis. These make a portable, cheap, and miniatured signal acquisition system. SPM has shown advantages in pathogens detection, such as H5N1, Zika virus, and SARS-CoV-2, due to its portability and high sensitivity (Ganguli et al. 2017; Yeo et al. 2016). Therefore, we build a portable SPM to detect the fluorescence triggered by RPA-CRISPR/Cas12a-virus recognition.

To detect the presence and concentrations of target pathogens, experts using professional software are required to analyze the results (von Chamier et al. 2021), which is time-consuming. Thus, we develop a novel method to accurately and quickly obtain the result from fluorescence images taken by SPM. Recently, machine learning and deep learning are used to quantify the concentrations of virus DNA from fluorescence images (Shiaelis et al. 2020), which makes the detection system integrated and time-consuming. Conventional neural networks (CNNs) are widely used to learn features from the raw pixelated images for a given classification task in an end-to-end manner (Liu et al. 2022; Lawrimore et al. 2019; Yang et al. 2021; Zhang et al. 2022). The most popular CNN-based deep learning models, such as AlexNet, DenseNet-121, and EfficientNet-B7, have been widely applied in medical image classification (Xie et al. 2020; Wang et al. 2021). Besides, in most cases, large data sets in specific domains are unavailable, which necessitates transfer learning (Artoni et al. 2020; Li et al. 2021). Transfer learning pretrains a deep learning model with a large dataset and the pre-trained model is used as the starting point for a new task with a small dataset, which can alleviate the demand for large data sets, overcome overfitting, and reduce training time (Yosinski et al. 2014). Herein, we use deep learning models with transfer learning for the classification of the fluorescence images.

In this study, we use a plate reader and achieve an LoD of 100 aM for purified MCP fragments with optimized RPA primers and crRNAs in 60 min. For POC detection, we build a portable SPM and achieve an LoD of 10 aM within 40 min. SPM outperforms commercial plate reader because of the systematic optimization of optical path and components. Besides, four animal-derived samples are used to confirm the reliability in practical application. To get results quickly and conveniently, we use three deep learning models, AlexNet, DenseNet-121, and EfficientNet-B7, with transfer learning to achieve accurate classification based on fluorescence images taken by smartphone microscopy. AlexNet performs well for binary (positive or negative, 100% accuracy) and multiclass (approximate concentration, 98.75% accuracy) classification. According to the results, we provide a fast, sensitive, easy-to-implement, and POC detection system for FV3.

## 2. Materials and Methods

### 2.1 Chemicals and reagents

We select 231 bp DNA fragments of the MCP gene from FV3 and Infectious spleen and kidney necrosis virus (ISKNV, *Iridoviridae*) as the target and control. Synthetic DNA fragments (FV3 MCP and ISKNV MCP) cloned into pUC57 vector, crRNA, and ssDNA-FQ reporter are purchased from Sangon Biotech. The sequences of the DNA fragments used in this work are shown in **Table 1**. PINDBK is purchased from Ebio company, Shenzhen, China. *Lachnospiraceae bacterium* Cas12a (LbCas12a) protein (M0653T) is purchased from New England Biolabs (Beijing) LTD. 96-well black microplate (3603) is bought from Corning Incorporated, New York, USA. UltraSYBR Mixture (CW0957M) is purchased from CWBIO, Taizhou, China.

### 2.2 Protein production and purification

UvsX, UvsY, GP32, and Bsu Plasmids are expressed in *E. coli* BL21 (DE3). The expression and purification procedures follow the published protocols (Yonesaki and Minagawa 1989). In brief, 0.5 mM IPTG is used to induce the protein expression. The incubation is at 16 °C with vigorous agitation (200 rpm) overnight for 20 hours. Cells are subsequently collected by centrifugation at 5600 g for 15 min at 4 °C. Pelleted cells are lysed via sonication (125 W, 15 min, work for 3 seconds, and pause for 8 seconds) in Lysis buffer (25 mM HEPES pH 7.5, 150 mM KCl, 10 mM imidazole, 1 mM TCEP, 0.01% Triton X-100, and 5% glycerol) supplemented with PMSF (1 mM), lysozyme, and DNase I. The lysate is clarified by centrifugation at 10000 g for 25 min at 4 °C. 1 mL of the Ni Sepharose High Performance (Cytiva) is added into the column and equilibrated with 5 to 10 column volumes of the Wash buffer (500 mM NaCl, 20 mM Tris-HCL, 1 mM DTT, 15 mM Imidazole, and 5% glycerol). Then, we load the pre-treated samples and wash them until the absorbance at 280 nm wavelength reaches the baseline (about 5 column volumes). Next, we elute the proteins with 20 to 30 mL Elution buffer. To obtain the suitable sample volume for next stage purification or enzymatic reaction, the eluted protein is pooled and concentrated to ∼40 mg/mL by using a 50 kDa cutoff ultrafiltration unit (Millipore). Finally, the purity of proteins is confirmed by SDS-PAGE (**Fig. S1C**).

### 2.3 Target amplification

We use a PCR amplification kit (Takara Bio, Japan) to obtain the purified dsDNA target and control with primers listed in **Table 1**. The PCR is run on a series multi-Block thermal Cycler PCR instrument (LongGene, China) with the following procedures: 95 °C for 5 minutes (1 cycle); 95 °C for 30 s, 60 °C for 45 s, and 72 °C for 30 s (35 cycles); 72 °C for 5 minutes (1 cycle); and 4 °C for storage. Agarose gel electrophoresis is used to confirm the DNA size, and dsDNA fragments are extracted and purified from the gel (Omega Biotek, D2500-02). The purified DNA fragments are quantified by using a Nanodrop 2000 spectrophotometer.

### 2.4 RPA reaction

We design RPA primers as described earlier (Liu et al. 2019). The sequences of primers are shown in **Table 1**. 50 μL RPA reaction contains 1 μL genome sample (with various concentrations), 0.2 μM forward and reverse primer, 16.2 μM UvsX, 5.5 μM UvsY, 7.2 μM Gp32, 4.5μM Bsu, and reaction buffer (TwistDx, Ltd., UK). Then, 100 mM MgCl_2_ is used to initiate the reaction. The assay is performed at 37 °C for 30 minutes (Huang et al. 2022).

### 2.5 CRISPR/Cas12a detection

We design crRNAs as described in our previous work (He et al. 2020a). LbCas12a-crRNA complexes are preassembled by incubating 1 μM LbCas12a and 1.25 μM crRNA at room temperature for 5 min. 1 μL RPA reaction product is dissolved in 1 × Binding Buffer (10 mM Tris-HCl, pH 7.5, 50 mM NaCl, 5 mM MgCl2, 0.1 mg/mL BSA), mixed with LbCas12a-crRNA complexes and 500 nM ssDNA reporter probe in a 100 μL reaction volume. The reactions are performed at room temperature for 30 minutes. The fluorescence signals are constantly examined by a microplate reader (SPARK, TECAN, Switzerland) at an excitation wavelength of 535 nm an emission wavelength of 595 nm, and a gain of 60. For POC detection, we perform the reactions at room temperature for 10 minutes, and the fluorescence signals are measured by SPM. To get a standard curve with SPM, various concentrations of purified targets are used, which include 100 pM, 1 pM, 10 fM, 1 fM, 100 aM, 10 aM, and 100 pM ISKNV MCP as control (before RPA). All data are normalized by negative control (divide the control value by the positive sample’s measurement), then integrated for two-sample *t*-test analysis.

### 2.6 SPM setup

We build the SPM by using a 532 nm green laser with 0.9 mW output power as the light source (Thorlabs, PL201), aspherical lens (Lubang), transmitted neutral density filters (Thorlabs, ND40A), dichroic mirror SEMROCK, FF555-Di03-25×36) with a cutoff wavelength of 535 nm (chrome, AT535), objective with 20x amplification (OLYMPUS OPLN20X), triplet achromatic lenses as the external lens (Thorlabs, TRH127-020-A), bandpass filter SEMROCK, FF01-542/27-25) and smartphone (Huawei Mate10). Translation stages, filter holders, and dichroic holders are bought from the Ruicage company. The instrument consists of two parts: the excitation path and the emission path. A laser beam passes through the neutral density (ND) filters to modulate the laser intensity. An aspherical lens (L1) away from the NFD generates a collimated beam. The collimated beam is reflected by the dichroic mirror (DM). The objective directs it onto the glass slide to illuminate and excite the sample forming the excitation path. The stage under the sample allows the fine adjustment of the focal plane, which directs the beam to the back focal plane of the objective (Vietz et al. 2019). Meanwhile, the emission beam is collected by the objective. On the other side of the objective, an external lens (L2) is positioned to form an intermediate image (Ning et al. 2021; Samacoits et al. 2021). The objective simultaneously illuminates the sample and collects the emission signal. Essentially the filter rejects excited light, allowing only the emitted light from the sample to reach the camera. The smartphone at the end of the emission path, set away from the external lens, records the fluorescence signal. The bandpass filter in front of the camera is used to optimize the detection of fluorescence signals. The glass slide and PDMS are treated as previously described (He et al. 2020b). The setup is immobilized on the breadboard for portable deployment.

### 2.7 Animal-derived samples

The animal-derived FV3 samples collected from wild or rearing animals are provided by the Animal and Plant Inspection and Quarantine Technical Center, Shenzhen, which include *Tiger frog virus* (TFV), *Bohle iridovirus* (BIV), *Soft-shelled turtle iridovirus* (STIV), *Rana grylio iridovirus* (RGV), and negative homologous sample. The information of the pathogens in this study is shown in **Table 2** (Zhao et al. 2007). We get the virus stored with preserving fluid (Biocomma, Shenzhen, China) and inactivated at 56 °C for 30 minutes. To release viral DNA, we add equal volumes of PINDBK to the samples and incubate them at 95 °C for 5 minutes. The numbers of the fluorescence images taken by SPM for the animal-derived samples are 14 for TFV, 15 for BIV, 24 for STIV, 25 for RGV, and 21 for negative sample.

### 2.8 Quantitative PCR reaction

The standard samples (purified target) and animal-derived samples are detected with the program: pre-denaturation at 95 °C for 10 minutes, followed by 45 cycles of denaturation at 95 °C for 10 seconds, annealing at 56 °C for 15 seconds, and extension at 72 °C for 30 seconds. Various concentrations (the highest concentration is 1 nM, the subsequent samples are diluted ten times each) of the purified target are used to get the standard curve **(Fig. S3B)**. We use PINDBK to get the viral nucleic acids from animal-derived samples and conduct the qPCR reaction with the same procedures.

### 2.9 Statistical analysis

Data are displayed as mean ± standard deviations (s.d.), with sample size indicated in the figure legend. The student’s two-sample unpaired *t*-test is used for statistical analysis. Statistical significance of the data is indicated as follows: *p < 0.05, **p < 0.01, ***p < 0.001, ****p < 0.0001; N.D. = no difference. GraphPad Prism 9 is used for statistical analysis.

### 2.10 Dataset and data augmentation

The datasets are fluorescence images collected in the detection assay for animal-derived samples and purified fragments **(Fig. 4A and S3A**). We use ImageJ to measure the mean grey value of each image. For data cleaning, we set an intensity range [median − standard deviation, median + standard deviation], if the images’ intensities are out of the range, they are considered outliers and excluded. After that, the number of the images for the purified target at certain concentrations is 8 for 100 pM, 9 for 1 pM, 10 for 10 fM, 1 fM, and 100 aM, 7 for 10 aM, and 20 for control. Increasing concentration of the purified target is respectively labeled with 0 – 6 in ascending order. The number of the images after data cleaning for the animal-derived samples are 12 for TFV, 11 for BIV, 23 for STIV, 22 for RGV, and 20 for negative sample, respectively labeled according to the concentrations determined by qPCR. Then, we get a dataset including 162 fluorescence images. Image augmentation including horizontal flipping, vertical flipping, and random noise is adopted before transfer learning-based deep learning to prevent overfitting and improve system robustness (Chlap et al. 2021). The number of fluorescence images is expanded from 162 to 337 for binary classification and 398 for multiclass classification.

### 2.11 Transfer learning

We adopt three deep learning models as our backbone network which includes AlexNet, DenseNet-121, and EfficientNet-B7 for binary classification and multiclass classification tasks. The input images are reshaped to 224 × 224 × 3 to satisfy the constraint of the pre-trained model, which is a common preprocessing for heterogeneous data in transfer learning. The training process has two steps. For the first stage of training, we leverage the weights of intermediate hidden layers derived from training and employ a pre-trained backbone network for feature extraction using the ImageNet dataset. Then, we replace the last fully connected layer having 1000 neurons for the ImageNet task with a fully connected layer consisting of 2 neurons or 7 neurons for the fluorescence classification task. For the second training, we fine-tune the full-scale backbone network with the fluorescence dataset and dynamically adjust the learning rate of the model to accelerate convergence via a function in PyTorch (Paszke et al. 2017). The performance of experiments is evaluated by using a series of metrics including confusion matrix, accuracy, precision, recall, and F1-score (Lawton and Viriri 2021).

## 3. Results

We extract the purified target and control DNA fragments that will be used for crRNA screening and LoD quantitation **(Fig. S1A)**. Amplification and collateral cleavage efficiency contribute to the sensitivity of detection. Thus, we design and test the RPA primers and crRNAs for the target DNA (Ding et al. 2021; Liu et al. 2022; Lu et al. 2020; Sasidharan et al. 2018). According to the preliminary results, the 4^th^ pair of RPA primers and crRNA-3 is the most efficient for amplification and collateral cleavage, respectively **(Fig. S1B and 2A)**. We perform the CRISPR/Cas12a reaction and achieve an LoD of 10 pM and 100 aM by crRNA-3 with and without RPA, respectively, which are higher than those by crRNA-1 and crRNA-2 (**Fig. 2B and 2C**). The integrated fluorescence signal of CRISPR/Cas12 and RPA-CRISPR/Cas12a by crRNA-3 is presented in **Fig.2D**. Using crRNA-3 and RPA 4^th^ primer, we decrease the LoD about 10^6^ times from 100 pM to 100 aM. Therefore, further experiments are performed using crRNA-3 and RPA 4^th^ primer.

**Fig. 1.**
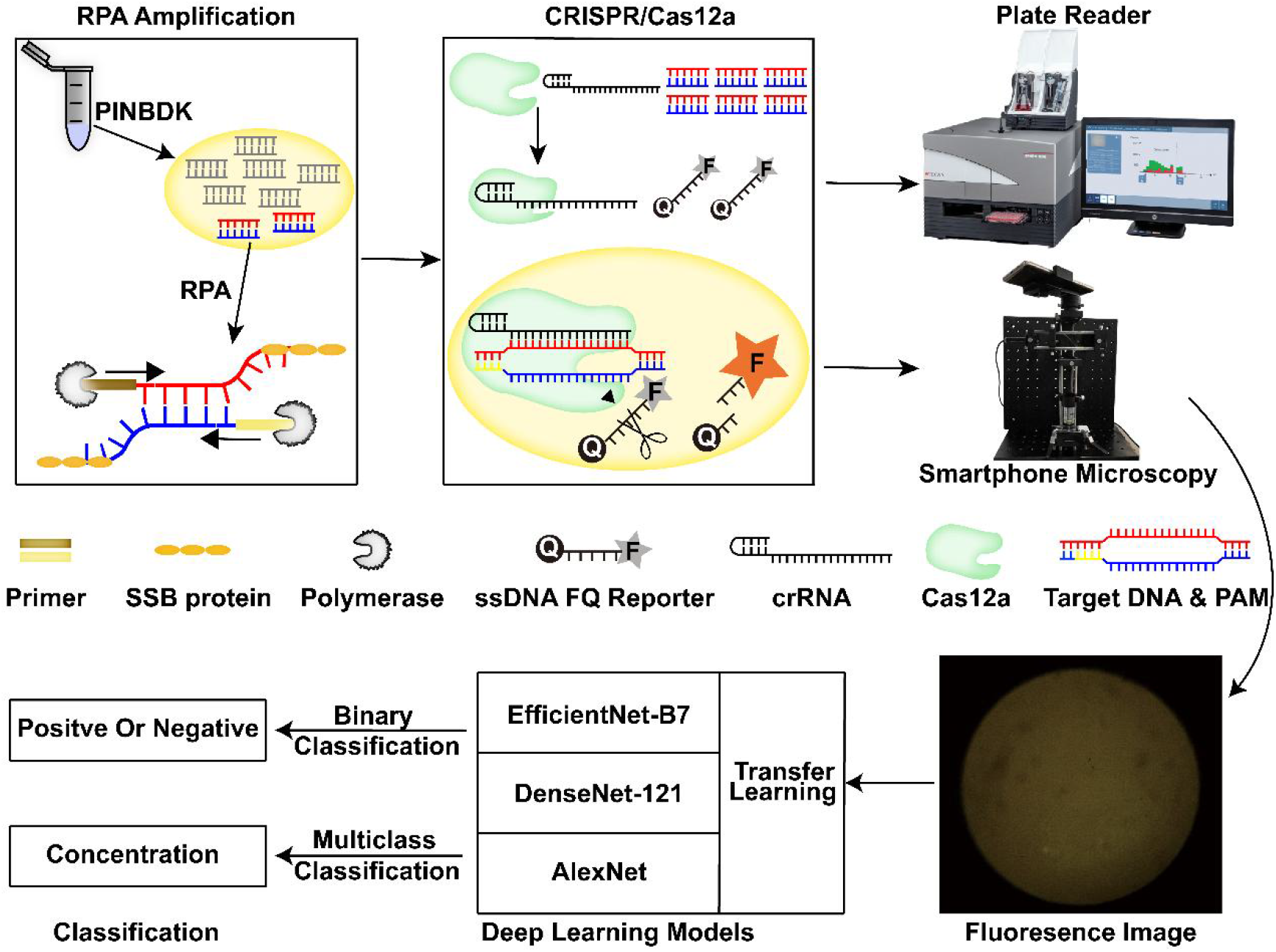
The schematic of the RPA-CRISPR/Cas12-SPM-AI detection system. The nucleic acids of animal-derived samples are released by Pathogen Inactive, Nucleic acid extraction, Direct-to PCR-Buffer with Proteinase K (PINDBK). Target DNA of the virus is amplified and recognized specifically by the RPA-CRISPR/Cas12a system. Cas12a bonds with crRNA and the Cas12a-crRNA complex bonds with target DNA, which triggers the collateral cleavage of Cas12a on the reporter (fluorophore quencher (FQ) labeled ssDNA probe). The fluorophore on the reporter is released and the fluorescence is detected by a commercialized plate reader or the SPM we build. Three different deep learning models including AlexNet, DenseNet-121, and EfficientNet-B7 with transfer learning are used to classify the fluorescence images.

**Fig. 2.**
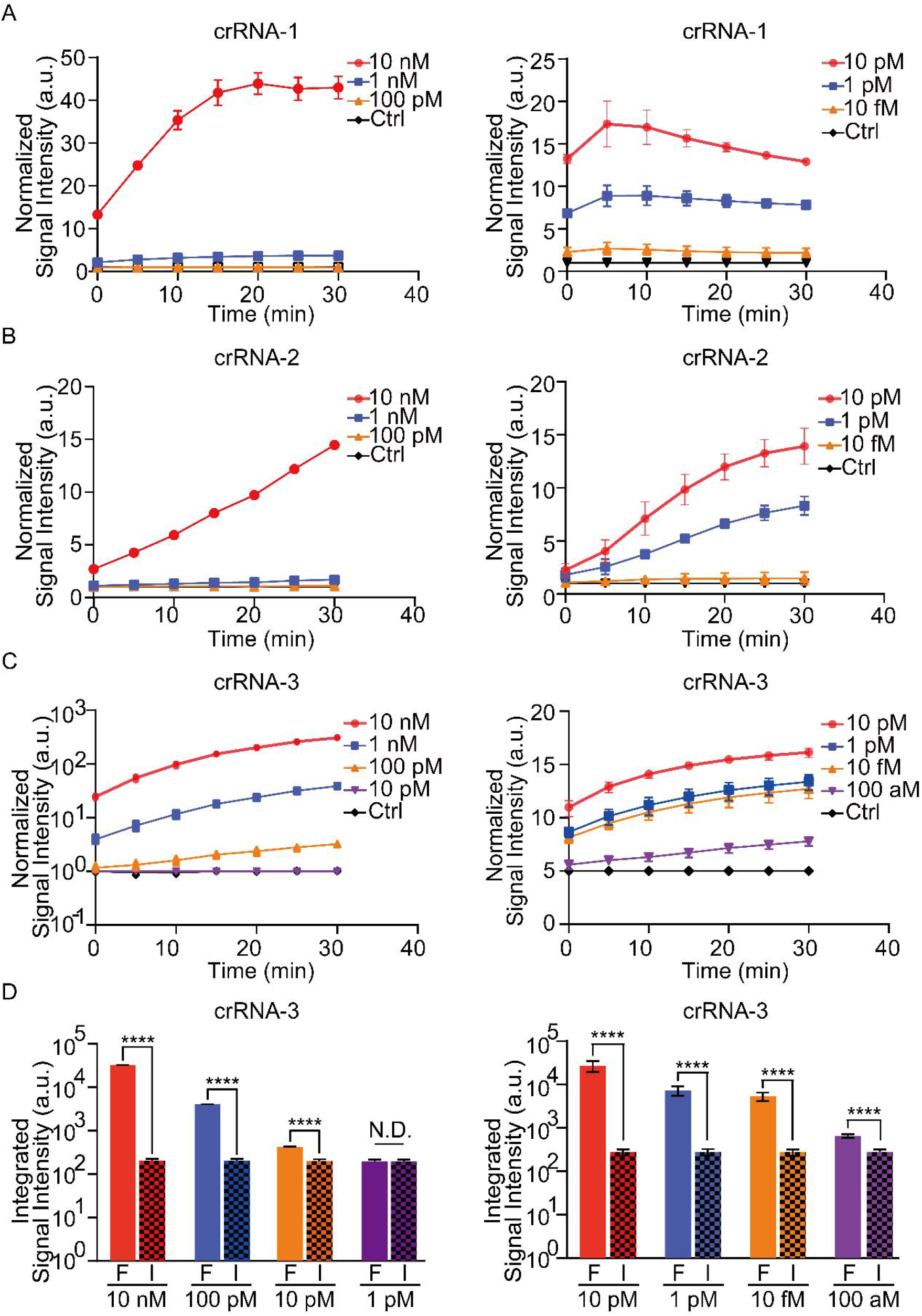
Detection of the purified fragments with three crRNAs by RPA-CRISPR/Cas12a system. **(A)** Normalized signal intensity of CRISPR/Cas12a (left) and RPA-CRISPR/Cas12a (right) by crRNA-1 for 10 nM to 100 pM target DNA versus 10 nM control DNA (left), and for 10 pM to 10 fM target DNA versus 10 pM control DNA (right), (incubation time: 30 min). (**B-C)** Similar experiments are performed by crRNA-2 and crRNA-3, respectively. **(D)** Integrated signal intensity of CRISPR/Cas12a (left) and RPA-CRISPR/Cas12a (right) by crRNA-3 for various concentration of target DNA versus 10 nM control DNA (left) and 10 pM control DNA (right), (incubation time: 30 min). F: FV3, I: ISKNV. Data are mean ± s.d. from at least three independent experiments. The student’s two-sample *t*-test is used for statistical analysis. *p < 0.05, **p < 0.01, ****p < 0.001. No error bar appears for certain points because it is shorter than the size of the symbol.

For POC detection, we build an SPM to detect the fluorescence signal generalized by the RPA-CISPR/Cas12a reaction. The schematic and physical view of SPM is shown in **Fig. 3A**. Standard samples are used to confirm the practicability of SPM (Chung et al. 2019). As shown in **Fig. S2A and S2B**, beads with a diameter of ∼1 μm and ∼4 μm can be observed, which indicates that the limit of resolution is approximately 1 μm. The images of the potato’s underground stem and stem tuber are shown in **Fig. S2C and S2D**. A significant change in signal intensity can be observed within 40 min including RPA for 30 min and CRISPR/Cas12a reaction for 10 min, as shown in **Fig. 2B**. Therefore, we decide to maintain the reaction time of CRISPR/Cas12a to 10 min for later experiments. To confirm the practicability of SPM in the detection system, we use it to quantitate the fluorescent signal triggered by RPA-CRISPR/Cas12a reaction. The fluorescence images are shown in **Fig. 3B**, which demonstrates that with the concentration of target DNA decreasing, the fluorescence signal is gradually reducing. This result is confirmed by ImageJ analysis (**Fig. 3C**). Thus, we achieve an LoD of 10 aM within 40 min. Besides, as shown in **Fig. S3A**, the fluorescence signal linearly increases with the elevated concentration of purified target DNA (Pearson’s R^2^ = 0.9404). Next, we detect five animal-derived samples including four positive and one negative samples. The fluorescence images and analyzing results are shown in **Fig. 3D and 3E**, which prove that animal-derived samples are detectable by the proposed system. To access the agreement between our system and qPCR, we compare the concentration of FV3 DNA from four positive samples using these two systems. The calculated concentrations of TFV, BIV, STIV, and RGV samples with our system are 4 pM, 59 fM, 1 fM, and 10 aM, respectively; 12 pM, 36 fM, 15 fM, and 45 aM with qPCR (**Fig. S3B**). The standard curve from the purified target and the fluorescence signal from the animal-derived samples are shown in **Fig. 3F**. As presented in **Fig. S3C**, The Pearson’s correlation coefficient is calculated to evaluate the correlation between qPCR and the proposed system (R^2^ = 0.9364, P < 0.0001). These results show a good consistency of our methods to the standard technique, which further demonstrates the reliability of our system.

**Fig. 3.**
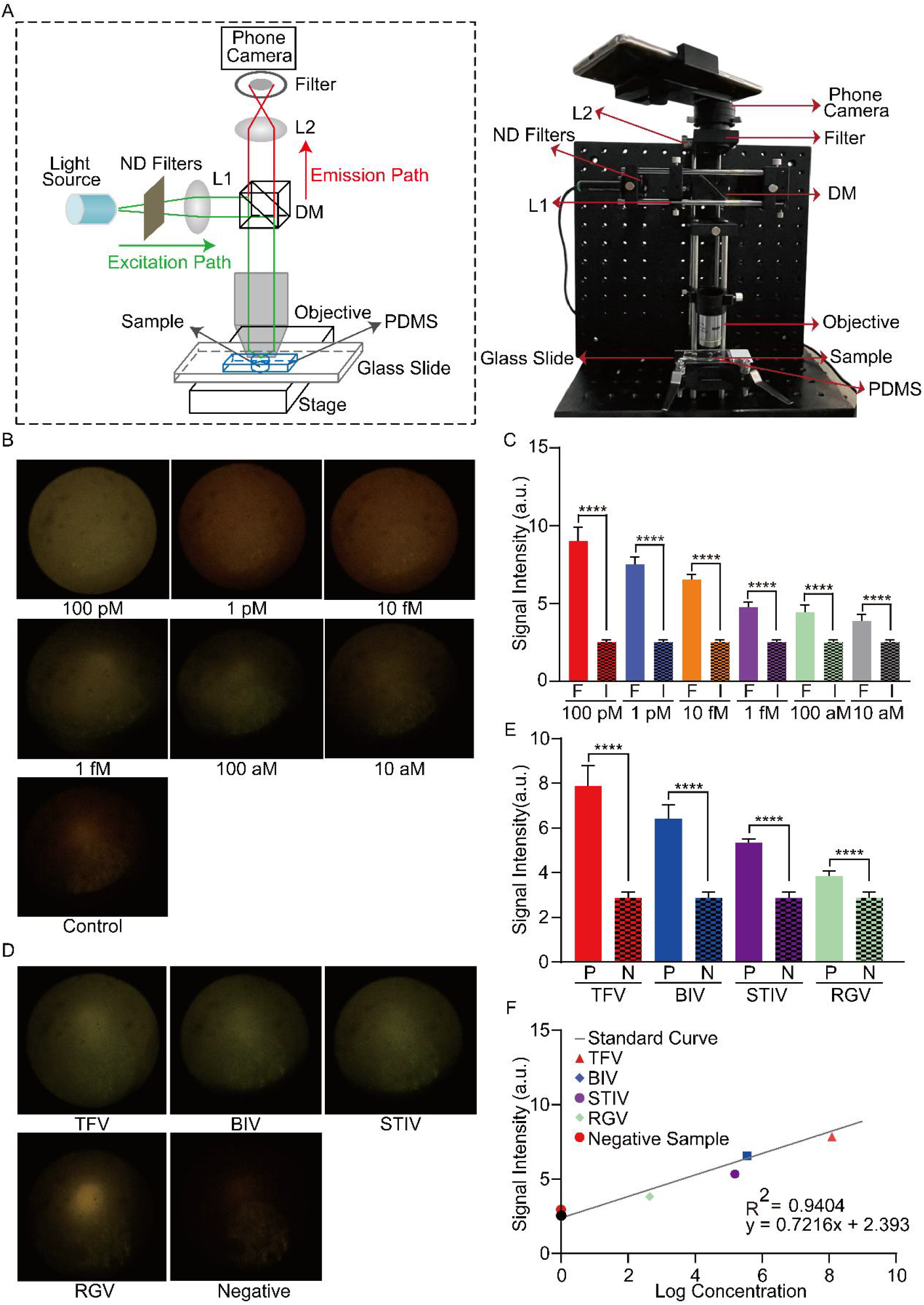
Detection of FV3 with RPA-CRISPR/Cas12a-SPM. **(A)** Schematic of SPM for fluorescence detection (left). The physical appearance of assembled device used for fluorescence image collection after the RPA-CRISPR/Cas12 reaction (right). DM: dichroic mirror, ND filters: neural density filters, PDMS: Poly-dimethylsiloxane. **(B)** The fluorescence images of purified target DNA and control DNA. **(C)** The statistics of signal intensity of fluorescence images from various concentrations of purified fragments. F: FV3, I: ISKNV. **(D)** The SPM images of the positive and negative animal-derived samples in the RPA-CRISPR/Cas12a detection assay. P: positive sample, N: negative sample. **(E)** The statistics of signal intensity of fluorescence images from animal-derived samples. P: positive, N: negative. **(F)** The fluorescence signal intensity of animal-derived samples and their absolute concentration detected by qPCR. The student’s two-sample *t*-test is used for statistical analysis. *p < 0.05, **p < 0.01, ***p < 0.001, ****p < 0.0001, N.D. indicates no difference.

The difference in fluorescence signal among positive and negative samples cannot be observed clearly with naked eyes **(Fig. 3B and 3D)**. Thus, to further improve the practicability and efficiency of this detection system, we use the three most advanced deep learning models with transfer learning to achieve the accurate classification of fluorescence images (C. et al. 2021; Hosny et al. 2020; Q. et al. 2014; Talo 2019). As shown in **Table 3**, Densenet-121, AlexNet, and EfficientNet-B7 all achieve decent performance with 100.00% F1 score, 100.00% recall, 100.00% precision, and 100.00% accuracy for binary classification in the positive or negative group. The accuracy and loss curve both reach a plateau, which indicates that there is no overfitting **(Fig. 4A and 4B)**. The confusion matrix shows that all test samples are correctly classified, which proves the reliability of the model **(Fig. 5C)**. Next, we evaluate whether the models can classify fluorescence images into multiple classes based on the concentration variation of target DNA. AlexNet gives the best performance, with 98.75% accuracy, 98.85% precision, 98.75% recall, 98.75% F1 score, and 15 ms inference time (**Table 4**). The confusion matrix shows that only two out of eighty test data are wrongly classified, which proves that we can get an approximate concentration by using transfer learning-based deep learning models without overfitting **(Fig. 4D-4F)**.

**Fig. 4.**
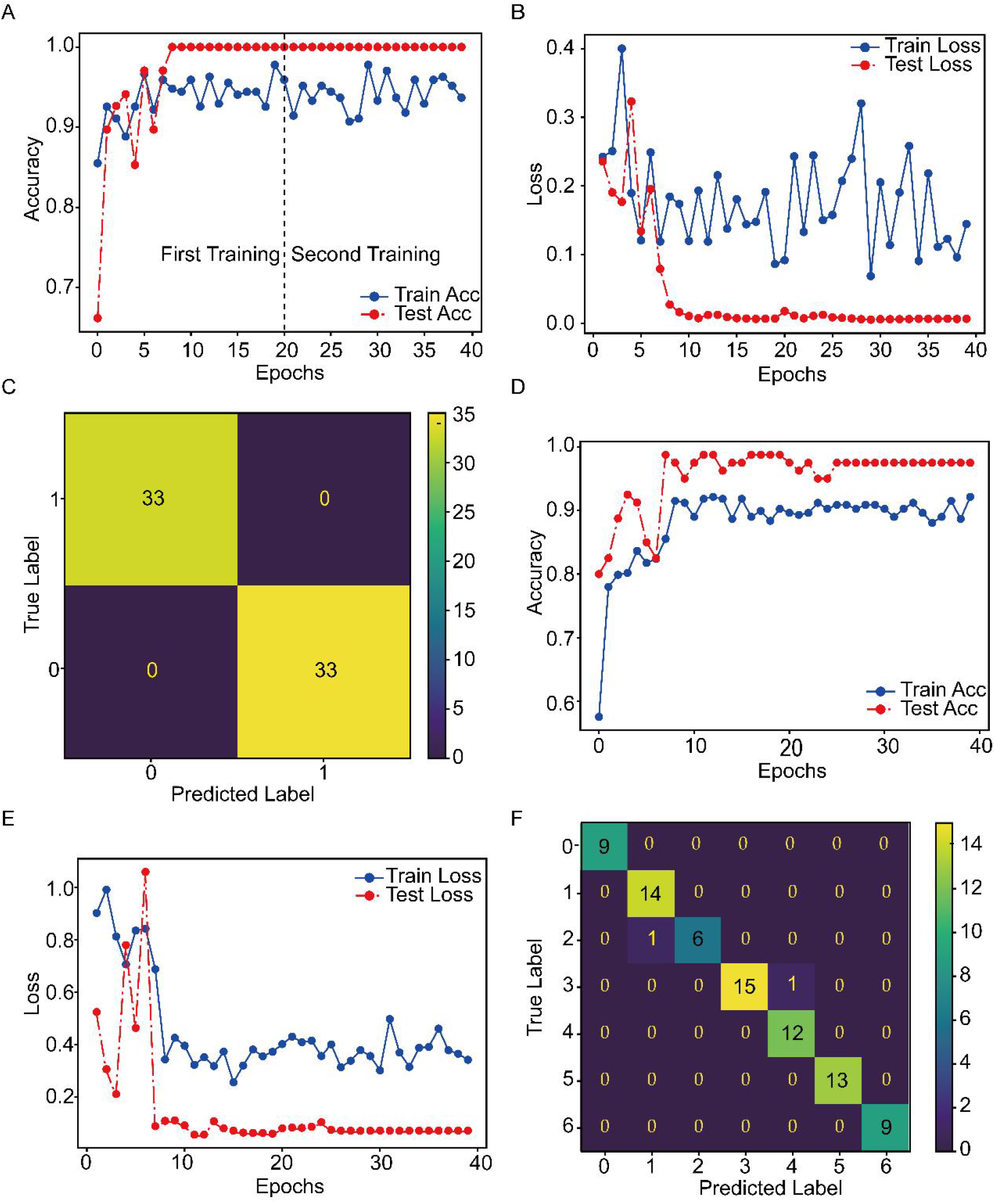
Evaluation of deep learning models with transfer learning for classification of fluorescence images. **(A-B)** Training traces of EfficientNet-B7 show the optimization of the accuracy and loss for binary classification. **(C)** Confusion matrix analysis of EfficientNet-B7 for binary classification. Each row of the matrix represents the instances in an actual class, while each column represents the instances in a predicted class. The diagonal in the matrix represents the ideal case in which the instance is correctly classified, while off-diagonal elements represent misclassified instances. **(D-E)** Training traces of AlexNet show the optimization of the accuracy and loss for multiclass classification. **(F)** Confusion matrix analysis of AlexNet for multiclass classification.

## 4. Discussion

In this work, we report the successful detection of FV3 using the RPA-CRISPR/Cas12a-SPM-AI system for the first time. We investigate from multiple perspectives to improve the specificity and sensitivity of the proposed system. The first approach is to design specific RPA primers and screen out the most efficient primers. The specificity of the primers is confirmed by blasting against the standard database of nucleotide sequencing, which can be used to amplify the target gene of the virus strains belonging to FV3. This result is also confirmed by detecting multiple kinds of animal-derived samples (**Fig. 3D and 3E**). However, recombination between FV3 and *common midwife toad virus* (CMTV, *Ranavirus, Iridoviridae*) is reported (Vilaca et al. 2019), which may result in cross detection, but it is very rare in nature. We believe that the minor flaw dose not limit the practicability of the system. Besides, different primers influence the amplification efficiency of the RPA reaction (Chen et al. 2020). However, there is no guide for designing the efficient primers, which requires the experimental validation (Fu et al. 2022). In this study, we design nine pairs of primers and find that the 4^th^ pair of primers show slightly stronger amplification efficiency among all candidate primers (**Fig. S1B**).

Collateral cleavage efficiency significantly contributes to the sensitivity of detection (Habimana et al. 2022; Liang et al. 2022). We design three crRNAs and find that crRNA-3 is the most efficient one. According to our results, we find that crRNA-3 triggers a stronger fluorescence signal without RPA. Besides, RPA-CRISPR/Cas12a decreases the LoD by crRNA-1, crRNA-2, and crRNA-3 about 10^4^, 10^4^, and 10^6^ times, separately **(Fig. 2)**. We speculate that it is because that the crRNA-3 recognizes the terminal of target DNA, while crRNA-1 or crRNA-2 binds with the middle region of target DNA. RPA is a method that the amplicons are produced from the end to the middle of the chain by consuming the raw materials such as ATP, dNTP, Mg^2+^, and active enzymes. (Lobato and O’Sullivan 2018). Exhaustion of the raw materials may generate amplicons that is not full length. This allows the region near the end of the target DNA to be amplified more efficiently and recognized by the crRNA. A similar result can be overserved from the other’s work (Chen et al. 2022). However, the opposite result is also reported (Jxab et al. 2022). This issue needs to be clarified with further experiments.

For POC detection, A portable SPM is built to detect the fluorescence signal. Fluorescence microscopy has been introduced for viral particle imaging and virus detection (Sivaraman et al. 2011; Wang et al. 2018). However, traditional microscopy is bulky and expensive for rapid POC diagnosis. Thus, the SPM setup is immobilized on the breadboard for portable deployment. Besides, it costs (approximately 2000 dollars) much less than a commercialized plate reader or real-time PCR machine.

An LoD of 10 aM is achieved with optimized RPA primers, crRNA, and SPM, which is comparable to the detection sensitivity of qPCR. FV3 DNA can be detected by PCR in the liver at 4 days post-infection (dpi) and in most organs at 14 dpi (Forzan et al. 2017b). The detection sensitivity of the proposed system is much higher than traditional PCR. Therefore, this optimized detection system could be deployed for early diagnosis with appropriate modifications. Besides, the collateral activity of the LbCas12a protein is constantly activated for 24 h (He et al. 2020b). We believe that the LoD can be further decreased by extending incubation time.

Compared to traditional detection methods and CRISPR-based detection system (Jiang et al. 2021; Ke et al. 2022; Kralik and Ricchi 2017; Shen et al. 2022; Tian et al. 2022), we obtain the result by deep learning-assisted classification within a very short period of time (15ms for multiclass classification of one image) without bulky instruments. Besides, transfer learning is adapted for deep learning models and a maximum of 98.75% accuracy for multiclass classification is achieved (**Table 4**). To the best of our knowledge, this is the first time that fluorescence images from RPA-CRISPR/Cas12a, taken by SPM are used to calculate the concentration of pathogen DNA with deep learning. Three deep learning models with transfer learning perform well on this task. We propose a possibility that classification based on fluorescence images does not require tiny features. The results indicate that a larger kernel of AlexNet (7×7) is more efficient on this task. In addition, to further optimize the detection system, we can deploy deep learning models on smartphones and get the result more conveniently and efficiently (Alkhulaifi et al. 2021; Sujit et al. 2021).

The development of detection methods for FV3 has been stagnant for many years. On the other hand, FV3 is becoming a global threat to biodiversity and aquaculture (Ferreira et al. 2021; Cozad et al. 2020; Oliveira et al. 2019). In this study, we establish, optimize, and evaluate the RPA-CRISPR/Cas12-AI system, providing a novel POC detection system for FV3. Furthermore, because the crRNA is reprogrammable, the integrated system could be used for detecting any DNA target. In terms of detection sensitivity, specificity, time, and reliability, the RPA-CRISPR/Cas12a-AI system will occupy an important place in the detection of DNA pathogens.

## 5. Conclusion

In this study, we develop an integrated, highly sensitive, easy-to-implement POC detection system for FV3. With the combination of RPA, CRISPR/Cas12a, smartphone microscopy, and deep learning-assisted classification, we achieve an LoD of 10 aM within 40 min. Without temperature regulation, this integrated system shows great potential for FV3 and DNA pathogens POC detection.

## Supporting information

Supporting materails

## Funding

This work is supported by the National Natural Science Foundation of China 31970752, Science, Technology, Innovation Commission of Shenzhen Municipality JCYJ20190809180003689, JSGG20200225150707332, JSGG20191129110812708, WDZC20200820173710001; Shenzhen Bay Laboratory Open Funding, SZBL2020090501004; China Postdoctoral Science Foundation 2020M680023; and General Administration of Customs of the People’s Republic of China 2021HK007

## Declaration of competing interest

The authors declare no competing interests that could have appeared to influence the work reported in this paper.

## Notes

### Competing Interest Statement

The authors have declared no competing interest.

