## Supporting materails for "Detection of Frog virus 3 via the system integrating RPA-CRISPR/Cas12a-SPM with deep learning"

**Title**

**Table 1** Oligonucleotides used in this study

| Ranavirus MCP | NTS: 5’ …gtaacccggctttcGGGCAGCAGTTTCGGTCGGCGTtcccaggtcg…3’ (231bp) |
| --- | --- |
|  | TS: 5’ …ccgacctgggaACGCCGACCGAAACTGCTGCCCtgctgcccgaaagc…3’ (231bp) ­­ |
| RPA primer F1 | ATGTCTTCTGTAACTGGTTCAGGTATCACAAG |
| RPA primer F2 | ATGTCTTCTGTAACTGGTTCAGGTATCACA |
| RPA primer F3 | TCTTCTGTAACTGGTTCAGGTATCACAAGTGGT |
| RPA primer R1 | GGCGTTGAGGATGTAATCCCCCGACCTGGG |
| RPA primer R2 | GGCGTTGAGGATGTAATCCCCCGACCTGGGAA |
| RPA primer R3 | CGTTGAGGATGTAATCCCCCGACCTGGGAACG |
| Ranavirus MCP F1 | ATGTCTTCTGTAACTGGTTCA |
| Ranavirus MCP F2 | GGCGTTGAGGATGTAATCCCC |
| ISKNV MCP F1 | ATGTCTGCAATCTCAGGTGC |
| ISKNV MCP F2 | GAGGTAGTCGCCGCCCCT |
| MCP qPCR F1 | GGTTCAGGTATCACAAGTGGT |
| MCP qPCR F2 | GCGTTGAGGATGTAATCCC |
| LbCas12a crRNA-1 | uaauuucuacuaaguguagauATCGACTTGGCCACTTATGACAA |
| LbCas12a crRNA-2 | uaauuucuacuaaguguagauTCAAGGAGCACTACCCCGTGGGG |
| LbCas12a crRNA-3 | uaauuucuacuaaguguagauGGGCAGCAGTTTTCGGTCGGCGT |
| ssDNA reporter | /5TAMRA/TTATT/3BHQ2 |

**Table 2** The information of animal-derived samples used in this study

| Pathogens | Host | Region | Refs |
| --- | --- | --- | --- |
| *Tiger frog virus* | Rana tigrina rugulosa | Guangdong,China | Unpublished |
| *Soft-shelled turtle iridovirus* | Trionyx Sinensis | Guangdong,China | Zhao et al. 2007 |
| *Bohle iridovirus* | Emydura krefftii | Guangdong,China | Unpublished |
| *Rana Grylio iridovirus* | Rana Grylio | Guangdong,China | Unpublished |

**Table 3** Deep learning models performance for binary classification

| Models  Index | DenseNet-121 | AlexNet | EfficientNet-B7 |
| --- | --- | --- | --- |
| Accuracy | 100.00% | 100.00% | 100.00% |
| Precision | 100.00% | 100.00% | 100.00% |
| Recall | 100.00% | 100.00% | 100.00% |
| F1 Score | 100.00% | 100.00% | 100.00% |
| Inference Time | 38ms | 24ms | 48ms |

**Table 4** Deep learning models performance for multiclass classification

| Models  Index | DenseNet-121 | AlexNet | EfficientNet-B7 |
| --- | --- | --- | --- |
| Accuracy | 97.50% | 98.75% | 97.50% |
| Precision | 97.67% | 98.85% | 97.66% |
| Recall | 97.50% | 98.75% | 97.50% |
| F1 Score | 97.48% | 98.75% | 97.47% |
| Inference Time | 31ms | 15ms | 41ms |

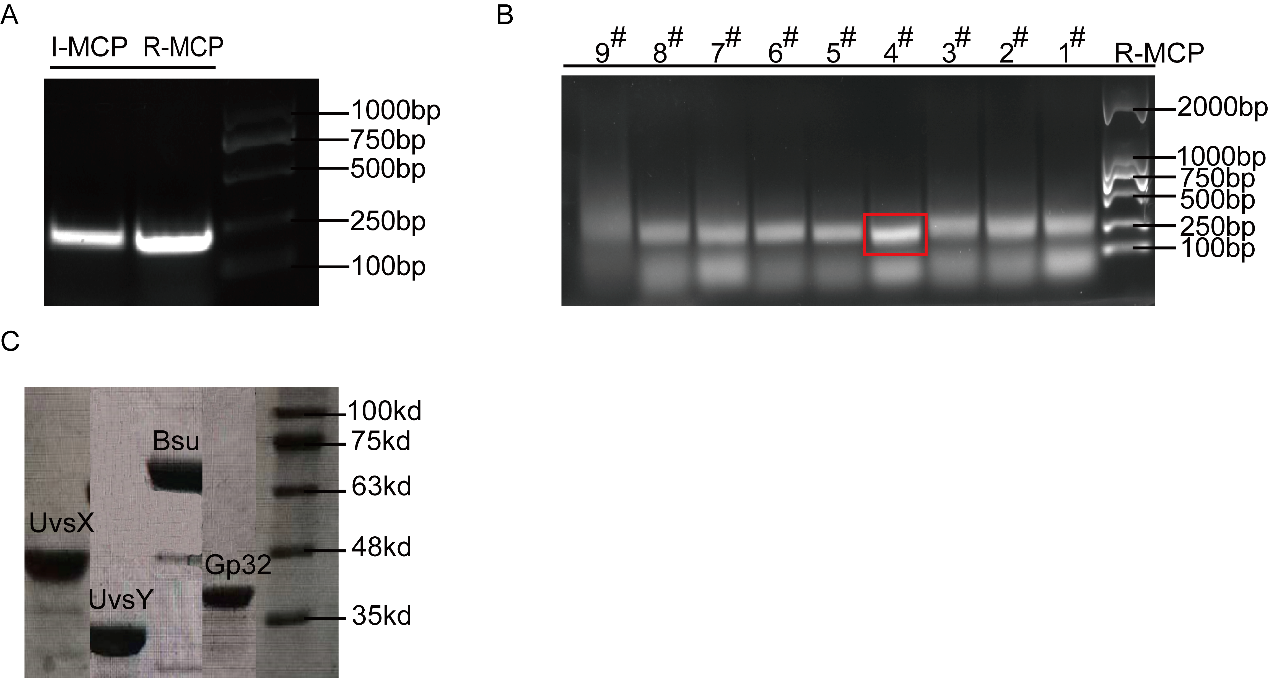

**Fig. S1.** Optimization of RPA primers and crRNA. **(A)** Agarose gel electrophoresis for purified target and control fragment. **(B)** Result of agarose gel electrophoresis for RPA efficiency. 1 nM purified target is used. 1^#^ represents RPA primer F1 and RPA primer R1. The results show that the 4^th^ pair of primers gives better amplification efficiency (RPA primer F2 and RPA primer R1). **(C)** The molecular weight of purified proteins for RPA is confirmed by SDS-PAGE.

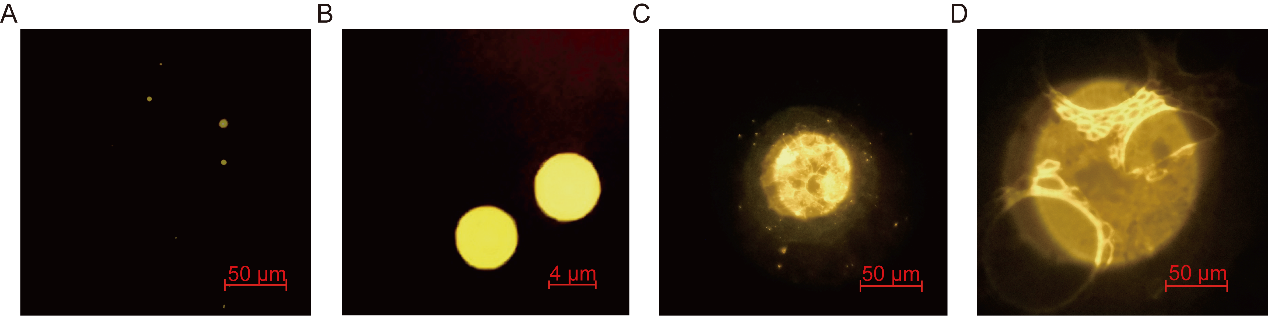

**Fig. S2.** Images of standard samples with SPM. **(A)** Detection of mixture beads with the diameter of 0.1μm, 0.2μm, 0.5μm, 1μm, and 4μm (TstraSpeck Fluorescent Microspheres Size Kit (T14792)). The excitation wavelength and the emission wavelength are 560 nm and 580 nm, respectively. **(B)** The image of 4 μm fluorescent beads. **(C)** The image of the potato’s underground stem. **(D)** The image of the potato’s stem tube

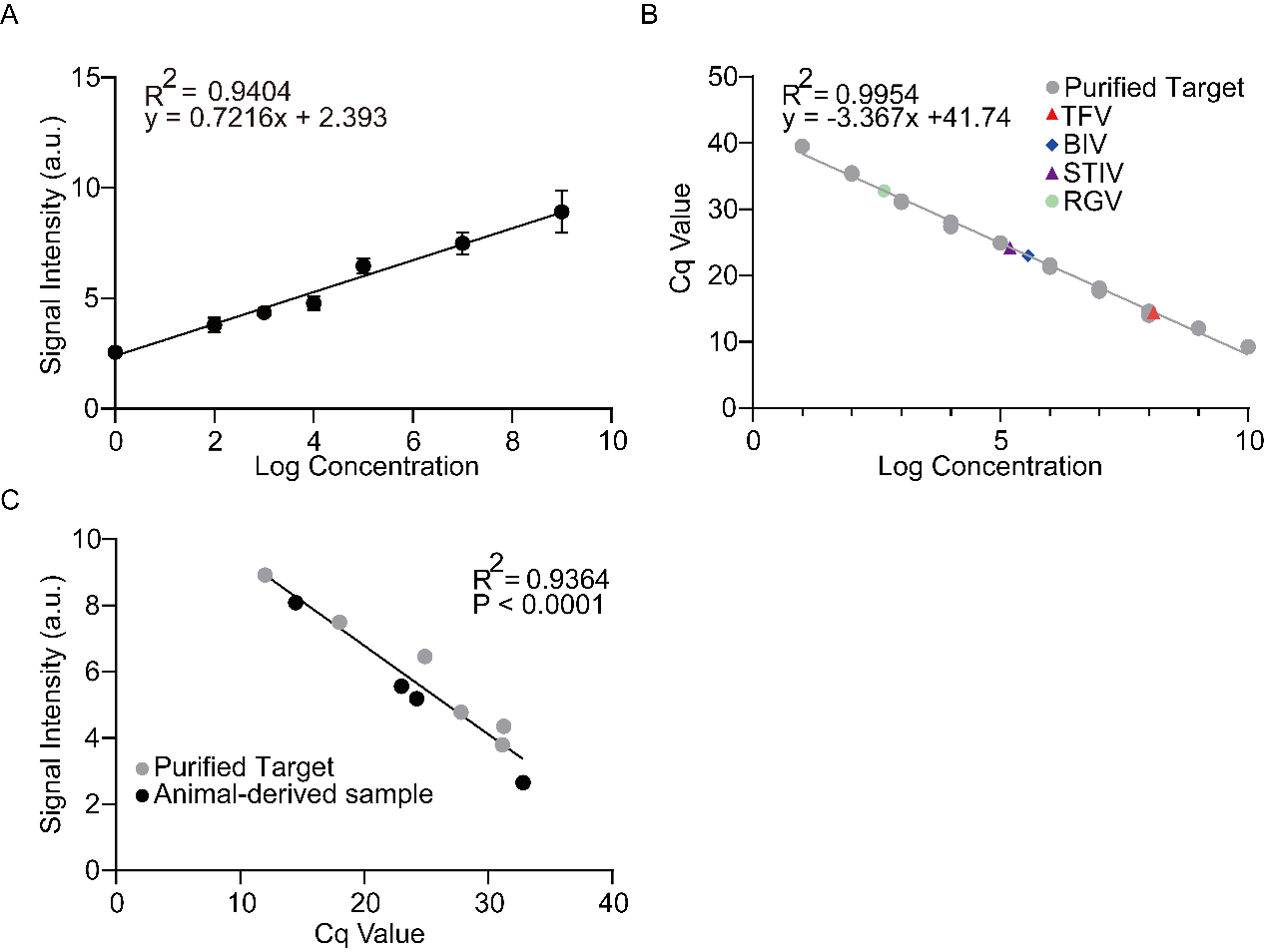

**Fig. S3.** Standard curve of the proposed system and qPCR. **(A)** Concentration-signal intensity curve of DNA concentration and smartphone images. The standard samples with purified target DNA of 100 pM, 1 pM, 10 fM, 1fM, 100 aM, 10 aM and control DNA of 100 pM are detected by crRNA-3 with RPA-CRISPR/Cas12a-smartphone microscopy**.** **(B)** Concentration-signal intensity curve of qPCR. The standard samples with purified target DNA of 1 nM, 100 pM, 10 pM, 1 pM 100 fM, 10 fM, 1 fM, 100 aM, 10 aM, and 1 aM are detected by qPCR, the concentration of each clinical sample is estimated according to this calibration curve. (C) The correlation analysis of FV3 detection results generated with the RPA-CRISPR/Cas12a-SPM system and commercial qPCR kit. The same sample detection Cq value of qPCR kit and the fluorescence signal intensity of the proposed system are co-analyzed.

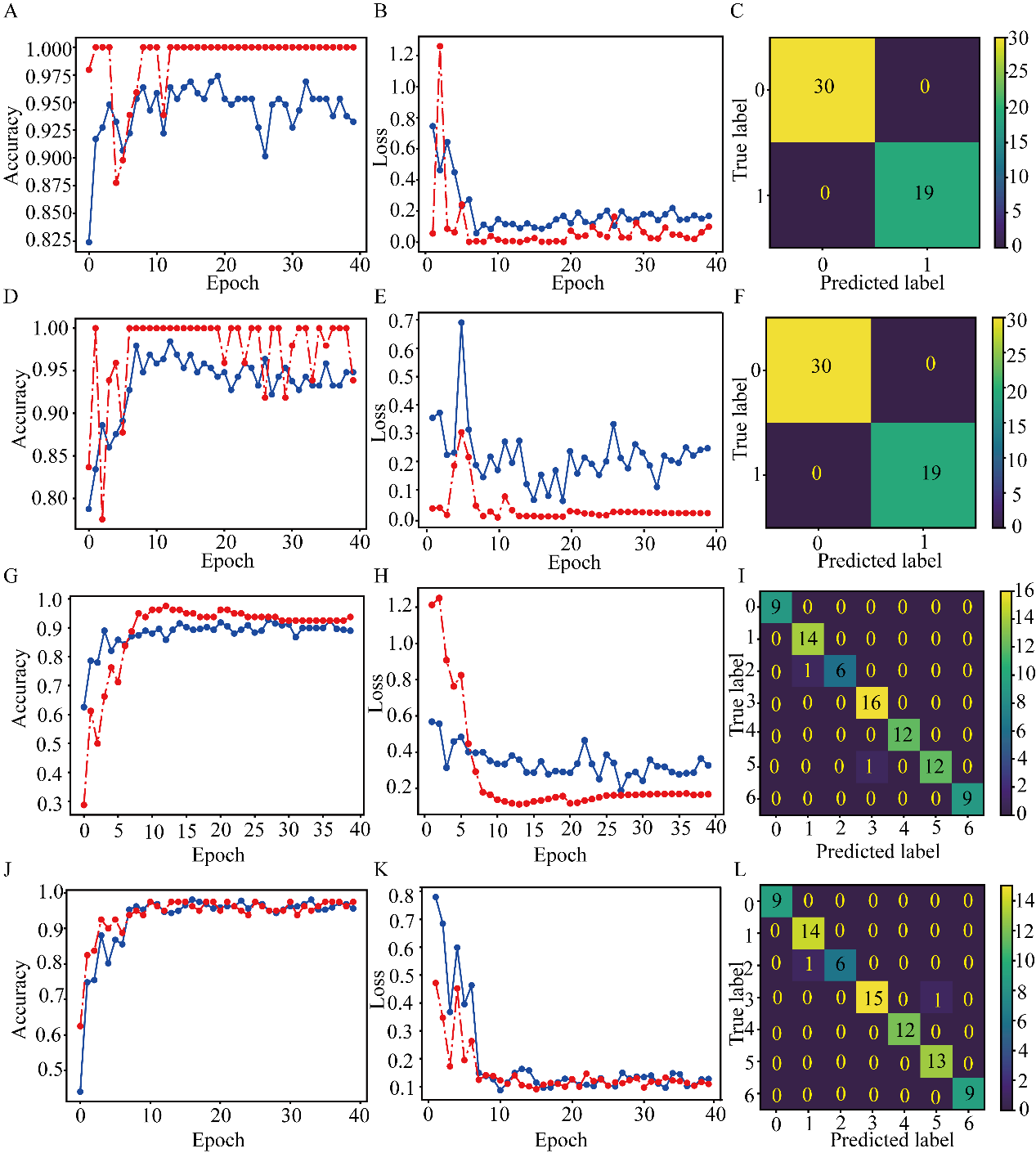

**Fig. S4.** Evaluation of Deep learning models for classification of fluorescence images. **(A)** Evaluation of DenseNet-121 for binary classification. **(B)** Evaluation of AlexNet for binary classification. **(C)** Evaluation of efficienctNet-B7 for multiclass classification. **(D)** Evaluation of DenseNet-121 for multiclass classification.
